# Microbial dark carbon fixation fueled by nitrate enrichment

**DOI:** 10.1101/2021.08.24.457596

**Authors:** Joseph H. Vineis, Ashley N. Bulseco, Jennifer L. Bowen

**Author notes:** Corresponding Author: Joseph H. Vineis, MSC(Marine Science Center), Nahant, MA 01908.

## Abstract

Anthropogenic nitrate amendment to coastal marine sediments can increase rates of heterotrophic mineralization and autotrophic dark carbon fixation (DCF). DCF may be favored in sediments where organic matter is biologically unavailable, leading to a microbial community supported by chemoautotrophy. Niche partitioning among DCF communities and adaptations for nitrate metabolism in coastal marine sediments remain poorly characterized, especially within salt marshes. We used genome-resolved metagenomics, phylogenetics, and comparative genomics to characterize the potential niche space, phylogenetic relationships, and adaptations important to microbial communities within nitrate enriched sediment. We found that nitrate enrichment of sediment from discrete depths between 0-25 cm supported both heterotrophs and chemoautotrophs that use sulfur oxidizing denitrification to drive the Calvin-Benson-Bassham (CBB) or reductive TCA (rTCA) DCF pathways. Phylogenetic reconstruction indicated that the nitrate enriched community represented a small fraction of the phylogenetic diversity contained in coastal marine environmental genomes, while pangenomics revealed close evolutionary and functional relationships with DCF microbes in other oligotrophic environments. These results indicate that DCF can support coastal marine microbial communities and should be carefully considered when estimating the impact of nitrate on carbon cycling in these critical habitats.

**Importance:** Salt marshes store carbon at one of the fastest rates of any blue carbon system and buffer coastal marine waters from eutrophication. Dark carbon fixation (DCF) conducted by microbes within the sediment can influence the carbon storage capacity, but little is known about the ecology or genomic potential of these organisms. Our study identifies a potential niche space for several functionally distinct groups of chemoautotrophs which primarily use sulfur oxidizing denitrification to fuel DCF under high nitrate concentrations. These findings fill an important gap in our understanding of microbial contributions to carbon storage within salt marsh sediments and how this critical blue carbon system responds to anthropogenic nitrate enrichment.

## Introduction

Salt marsh sediments store carbon at a higher rate than any other coastal marine habitat (1) and the accumulation of carbon is influenced by diverse heterotrophic and autotrophic microbial engineers inhabiting the sediment. Macrophyte communities dominate the surface of the salt marsh platform and contribute significant amounts of carbon in the form of roots, shoots, and exudates (2,3). Sulfate reduction is the primary respiratory process used by the heterotrophic microbial community to decompose macrophyte derived carbon sources (4), resulting in a loss of photosynthetically produced carbon and the production of large pools of reduced sulfur and ammonium. These pools of reduced compounds in turn can be used by chemoautotrophic microbes to fix inorganic carbon in the absence of light using dark carbon fixation (DCF) (5–7). Near-shore and shelf sediments are among the most important contributors to oceanic carbon fixation (0.29 Pg C yr ^-1^) (8) and primary production (9). Thus, chemoautotrophic microorganisms could represent a significant contribution to carbon storage in salt marsh sediments, based on evidence from molecular genomics (10), carbon isotopes (10,11), and relevance in other marine systems (8,12).

Many chemoautotrophs inhabiting marine sediments primarily use either the Calvin Benson Bassham (CBB) or the reductive TCA (rTCA) cycle to fix inorganic carbon (7) using energy derived from the oxidation of reduced sulfur or ammonium (7). Facultative sulfide oxidizing bacteria (SOB) can generate energy for DCF using the multienzyme sox pathway or through a reverse-acting dissimilatory sulfite reductase (rDSR) coupled with an adenosine 5’-phosphosulfate (APS) reductase and an ATP-sulfurylase that completely oxidizes reduced sulfur to sulfate (5,13–16). Nitrate can be used as the terminal electron acceptor for the oxidation of reduced sulfur in SOB within marine sediments and increased nitrate availability can simultaneously stimulate production of sulfate and nitrous oxide (17). Current knowledge of the physiology, phylogeny, and niche space of microbial populations that govern DCF is limited, hampering our ability to accurately predict how anthropogenically driven changes in climate and biogeochemistry will influence carbon storage in coastal marine sediments. Examining the genomic capacity of microbes in salt marsh sediments exposed to high concentrations of nitrate will provide important insight on the relative importance of autotrophic vs. heterotrophic nitrate reduction in coastal systems exposed to excess nutrients.

To improve our understanding of microbial contributions to carbon cycling within coastal sediments, we reconstructed draft genomes from metagenomic data (MAGs) derived from a previous NO_3_^-^ enrichment experiment (18). The recovery of draft genomes allows us to resolve several outstanding unknowns regarding DCF within salt marsh sediments including: 1) Common metabolic strategies and traits potentially important to defining the niche space; 2) The vertical niche space boundaries among chemoautotrophs; and 3) phylogenetic and functional relationships with other chemoautotrophs. To address the first two unknowns, we characterized MAGs across a sediment depth profile and identified genes related to DCF, decomposition of organic carbon, and sulfur and nitrogen cycling. To address the third unknown, we used a pangenomic approach to identify genes that are potentially important to inhabiting coastal sediments.

## Methods

### Sampling from flow through reactors and the environment

We collected 27 sediment samples from a flow through reactor (FTR) experiment described previously (18,19), where every effort was made to prevent the influence of light and oxygen. We collected three replicate sediment cores from the tall ecotype of *Spartina alterniflora*.Sediment from “shallow” (0-5 cm), “mid” (10-15 cm), and “deep” (20-25 cm) regions from each core were sectioned and homogenized independently under anoxic conditions and placed into two separate FTRs (n = 18). Nine FTRs received 500 μM K^15^NO_3_^-^ in sterile sea water (“N-enriched”) and the other nine received sterile sea-water alone (“unenriched”) at a flow rate of 0.08 mL min^-1^. After 92 days, we collected sediment from each of the reactors. Sediment was also collected from the homogenized depth sections of each core prior to the beginning of the FTR experiment (“pre-enriched”). An additional 6 samples were collected from the top 5 cm of sediment from two creeks located in the same marsh, part of the Plum Island Long Term Ecological Research (PIE-LTER) marsh complex (“in-situ”). The in-situ samples were collected in May, July, and September from a N-enriched and a paired reference creek area dominated by *Spartina alterniflora* and described in Kearns et al. (2016). Additional metadata can be found in Table S1.

### MAG reconstruction and quality evaluation

Details of metagenomic DNA extraction, sequencing library preparation, quality filtering, and metagenome assembly are included in supplemental methods. We reconstructed draft genomes for each assembly and summarized the quality of each metagenomic assembled genome (MAG) using Anvi’o v5 (20). We do not report the assembly statistics or MAGs from the in-situ samples due to a highly fragmented assembly.

### Identification of carbon fixation pathways and nitrogen, sulfur, and organic carbon cycling potential among MAGs

Functional annotation of genes related to carbon, nitrogen, and sulfur cycling was carried out using DRAM (21). Confirmation of DRAM annotation and genome queries of additional functional genes were conducted using HMM models obtained from Fungene (22) and the Carbohydrate Active Enzymes database (CAZy) (23). The search for additional functional genes using Fungene and CAZy HMM models was conducted using HMMER and facilitated by Anvi’o. Metabolic potential was also estimated for each of the genomes according to the RAST genome annotation tool (24). We estimated the percentage of genes in two well-known carbon fixation pathways, the Calvin-Bensen-Bassham (CBB) and reductive TCA (rTCA) cycle, according to the detection of several genes in each of the pathways during DRAM annotation and confirmed using RAST. We used phylogenetic confirmation to ensure the accuracy of assigning CBB and rTCA to each MAG (Supplemental). The HMM models we used and a table containing gene names and links to HMM models are reported in Table S2 and custom Anvi’o HMMs are available here (https://github.com/jvineis/Reactor-metagen/).

### Phylogenetics of estuary environmental genomes

We built a phylogenetic tree of FTR MAGs and all MAGs in the Genomes from Earth’s Microbiomes (GEM) (25) database that were recovered from marine intertidal and salt marsh habitats to identify evolutionary relationships among the FTR MAGs and other coastal environments. Details of the MAGs, samples included, and the methods employed can be found here https://github.com/jvineis/Reactor-metagen/.

### MAG taxonomic classification, dereplication and relative abundance

We used the GTDB-Tk classifier (26) to assign a taxonomic classification to each MAG (GTDB) (27). Average nucleotide identity (ANI) among all MAGs was computed using pyani (28) employed with Anvi’o v7. MAGs with an ANI of 95% over 90% of their genome were de-replicated to estimate their relative abundance (RA) in each of the samples. The contigs of the dereplicated collection of MAGs were concatenated into a single collection and each short-read dataset was mapped to this collection using bbmap (29). We used samtools (30) to filter, sort, and index the mapping results. MAG relative abundance in each sample was calculated to allow for comparison of relative abundance across and within samples according to the following equation:

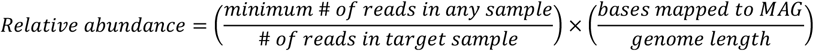

We estimated the occurrence of each MAG within each experimental group by dividing the sum of the relative abundance of a MAG within an experimental group by the total relative abundance observed across all experimental groups.

### Pangenomics of Chlorobium and Sedimenticola

We used reference genomes for a pangenomic analysis to identify functional and evolutionary relationships among the chemoautotrophs we recovered. We selected *Chlorobium* and *Sedimenticola* as representative chemoautotrophs that employ rTCA and CBB respectively, because we assembled multiple *Chlorobium* and *Sedimenticola* MAGs. There are also several well studied reference genomes available for comparison. We obtained 20 *Chlorobium* and 14 *Sedimenticola* reference genomes from IMG and NCBI with completion estimates above 50% and redundancy below 10% according to the collection of single copy genes used by Anvi’o v7. We employed a pangenomic analysis, including the identification of gene clusters, and ANI among the MAGs and reference collections using Anvio v7.

### Data availability and reproducibility

We provide open access to each of the bioinformatic steps here: https://github.com/jvineis/Reactor-metagen/. The files required for visualization of contig coverage across samples and placement into respective MAGs are located here: https://figshare.com/articles/dataset/Anvio_Files/12034611. Raw sequence reads are archived on the MG-RAST server under Project Numbers mgp84173 and mpg99679.

## Results

### MAG reconstruction

Genome reconstruction of the 27 metagenomic samples from the FTR experiment produced a collection of 113 MAGs that were greater than 48% complete and less than 7% redundant according to a collection of 71 single copy genes, with average completion and contamination estimates of 79.0% and 1.8% respectively. With the exception of assembled rRNA genes (Table S3), 44 of the MAGs met all the criteria of high-quality MAGs (31). Mean coverage estimates for the high-quality MAGs, based on mapping reads used to assemble the MAGs, ranged from 5.7 – 53.9 with an average of 15.5 (Table S1). The number of MAGs recovered from each of the samples ranged from 0-15 (Table S1) and the vast majority (93 of 113) were recovered from the N-enriched samples. Reconstruction of MAGs was most successful from mid depth and deep samples, as opposed to shallow samples, with a total of 33 and 42 MAGs, respectively. The percentage of the metagenome mapping back to our collection of MAGs was also highest in N-enriched samples, where read recruitment ranged from 6.5 – 27.3%, with a mean of 13.0%. We did not observe more than 2.4% read recruitment in either the pre, unenriched, or in-situ samples (Table S1).

### Chemoautotrophic functional groups, sulfur oxidation, and denitrification

We assessed chemoautotrophy using CBB and rTCA within each MAG using aclA/aclB and cbbS/cbbL respectively and phylogenetic confirmation of the genes for each MAG are shown in Fig. S1. To provide additional evidence for chemoautotrophy in each MAG, we calculated the proportion of the 10 and 9 genes assessed for pathway completion in the rTCA and CBB pathways respectively (Fig. 1, Fig. S1, Table S2). We did not place a threshold on the number of genes required to be considered a competent chemoautotroph and instead relied on the presence of *aclA/aclB* and *cbbS/cbbL* (32,33). According to the presence of these genes, we found evidence of inorganic carbon fixation potential coupled with denitrification and sulfur oxidation in 59 of 113 MAGs (Fig. 1). Genomic potential indicative of denitrifying sulfur oxidation was detected within five distinct functional groups of chemoautotrophic MAGs (Fig. 1.).

**Figure 1.**
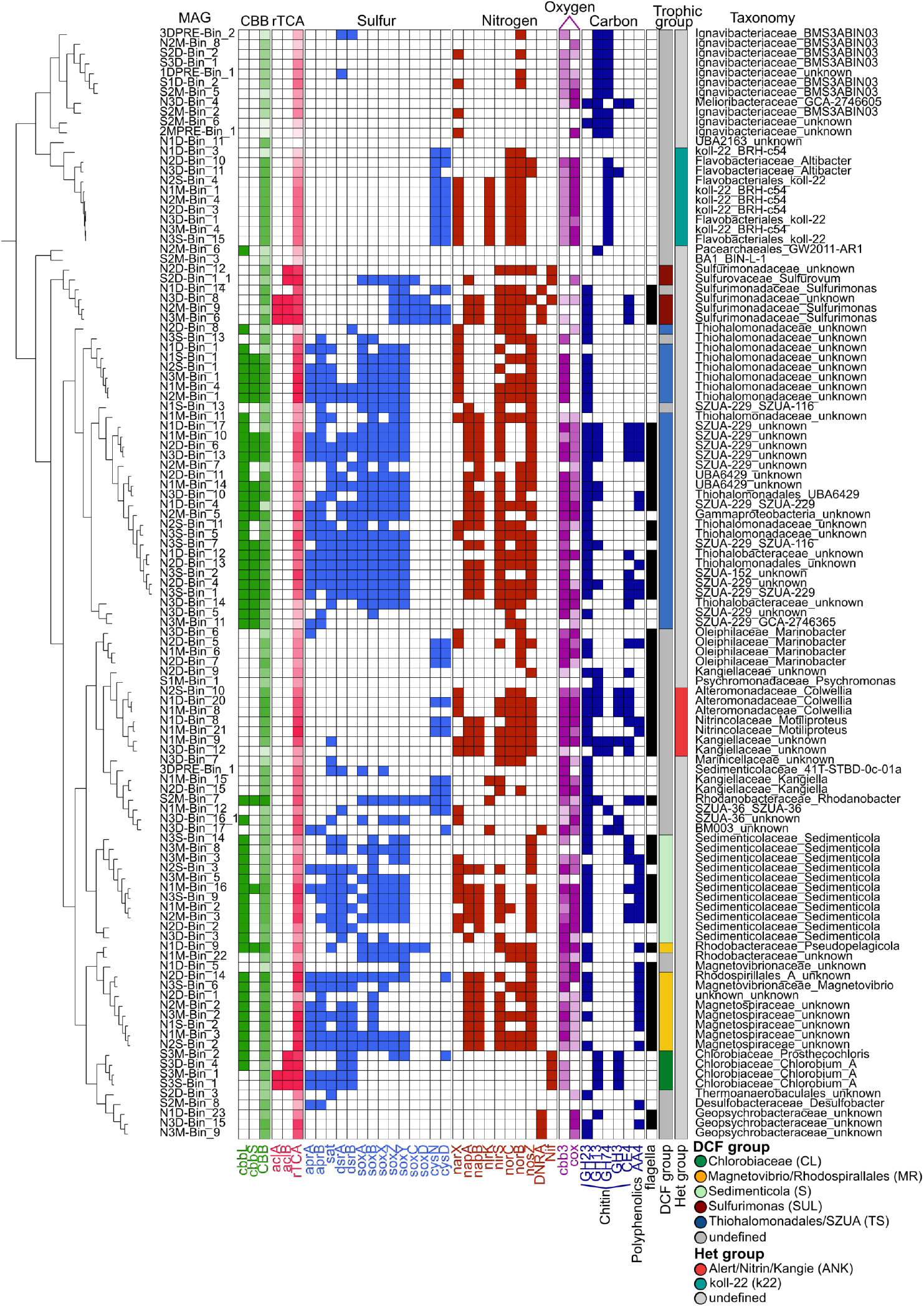
Functional description of all MAGs recovered. From left to right 1) Functional clustering: the figure displays a hierarchical clustering of genomes based on a presence absence matrix of all functions identified by RAST. 2) The unique name for each MAG. The MAG naming convention includes the source of the MAG (N = enriched, S = unenriched), the core ID (1, 2, or 3), the sediment depth (S= shallow, M= mid, D = deep), the MAG (Bin) ID. A heat map displaying the detection of genes for each specific target function. The genes searched for each function are reported in Table S2. The functions in the heatmap are colored based on genes in each functional category: green = Calvin-Benson-Bassham, pink = reductive TCA cycle, blue = sulfur cycle, red = nitrogen cycle, dark blue = select carbohydrate utilization genes, and black = flagella. The variable heat displayed in the CBB and rTCA columns of the display represent the percentage of 9 and 10 genes used to evaluate the complete pathway respectively and can be found in Table S2. The functional categories for MAGs (Trophic group) classified as functionally capable of dark carbon fixation are displayed in the DCF group column and the heterotrophic canonical denitrifier groups are displayed in the Het group column. The functional groups in the legend for dark carbon fixation (DCF group) include Chlorobiaceae (CL), Magnetovibrio/Rhodospirallales (MR), Sedimenticola (S), Sulfurimonas (SUL), Thiohalomonadales/SZUA (TS) and undefined. The (Het group) includes two main clusters of MAGs, the Alteromonadacea/Nitrincolaceae/Kangiellacaea (ANK), and the koll-22 (k2). The GTDB-Tk taxonomy are presented in the last column.

MAGs of three of the functional groups (*Thiohalomonadales/SZUA* (TS), *Sedimenticola* (S), and *Magnetovibrio/Rhodspirallales* (MR)) contained the potential for CBB, according to the presence of *cbbL* or *cbbS* (Fig. 1). These groups could be broadly classified as denitrifying sulfur oxidizers, containing genetic evidence for at least one marker gene in the denitrification pathway: nitrate reductase (*narGHI/napAB*), nitrite reductase (*nirSK*), nitric oxide reductase (*norCB*), and/or nitrous oxide reductase (*nosZ*) (Table S4). We identified complete denitrification potential in 17 MAGs identified with CBB potential in these three functional groups (Table S4). Sulfur oxidation potential of the MAGs within these three CBB functional groups was commonly characterized by the presence of the SOX gene complex (*soxABXYZ*), dissimilatory sulfite reduction (*dsrAB*), APS reductase (*aprAB*), and ATP sulfurylase (*sat*). We graphically characterized the co-occurrence of these genes for a *Sedimenticola* MAG in the S group to demonstrate the interconnectivity of the sulfur, nitrogen and carbon related metabolic potential (Fig. 2).

**Figure 2.**
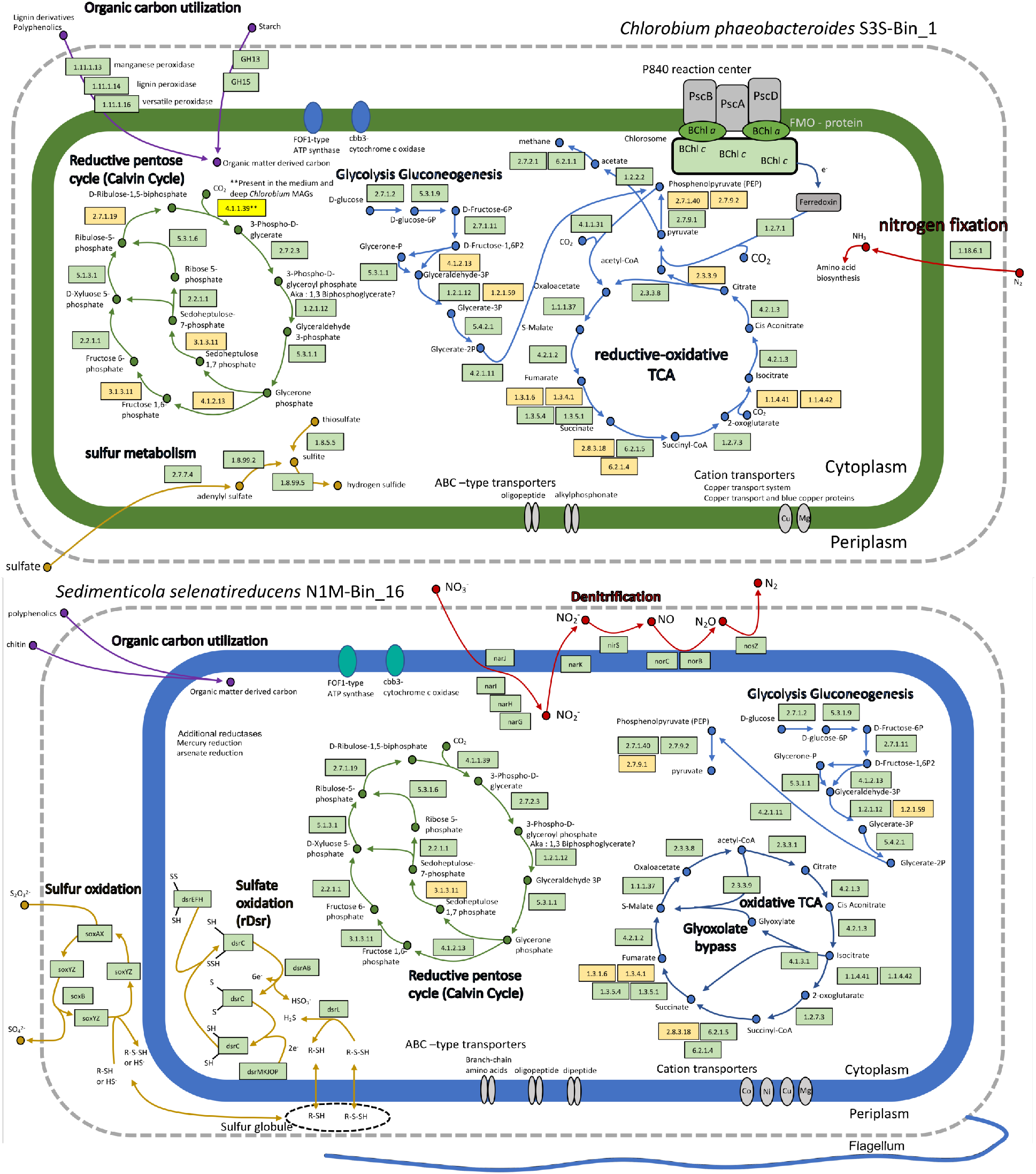
A graphical summary of the functional potential of an A) *Sedimenticola* and B) *Chlorobium* MAG based on RAST and DRAM annotation. The *Sedimenticola* MAG represents a member of the TS group (Fig. 1) which uses the CBB carbon fixation pathway powered by denitrifying sulfur oxidation. Each reaction is indicated by an enzyme commission (EC) number or gene name within a green or yellow box. Green shading of the EC numbers indicates the presence of the gene in our MAG and a yellow background indicates the gene is absent. The substrates and products are indicated by a filled circle. Pathways include the potential primary carbon fixation pathway, metabolism of sulfur, nitrogen and key central carbon metabolisms of the cell.

The *Sulfurimonas/Sulfurimonadaceae* group (SUL) members contained genomic potential for carbon fixation via the rTCA cycle, sulfur oxidation via a *soxCDYZ* complex, and sulfate assimilation through sulfate adenylyltransferase (*cysDN*) (Fig. 1). The SUL group also contained genes for either dissimilatory nitrate reduction to ammonia (*nrfAB*) or nitrogen fixation (*nifH*) in addition to at least one gene in the denitrification suite (Table S4). We detected evidence of the rTCA cycle among the *Chlorobiaceae* group (CL), which also contained genes for dissimilatory sulfate reduction (*dsrAB*) and (*aprAB*) and nitrogen fixation (*nifH*). Based on the genomic characterization of the CL group, they are likely photoautotrophs with the potential capacity for mixotrophic growth. We graphically characterized gene pathways for a representative *Chlorobium* MAG demonstrating this mixotrophic capacity (Fig. 2).

MAGs representing each of the four chemoautotrophic functional groups contained the potential for chitin utilization and polyphenolics (Fig. 1). Additionally, electron transport complexes were detected within these MAGs, including oxygen dependent cytochrome c oxidase, cbb3 type (cbb3), and the cytochrome c oxidase, type IV (cox) (Fig.1). We also observed the potential to produce flagella in both heterotrophic and DCF functional groups except for the CL and k22 groups (Fig. 1).

Canonical heterotrophic denitrifiers were also recovered from each of the N-enriched FTR sediments, including seven complete denitrifiers (Fig. 1, Table S4). The canonical denitrifiers formed two main functionally similar groups (k22, ANK), with the first (k22) containing MAGs with taxonomic classification as “koll-22” and a second containing MAGs with *Alteromonadaceae*, *Nitrincolaceae*, and *Kangiellaceae* classification (ANK) (Fig. 1). Many partial and complete heterotrophic denitrifiers also contained assimilatory sulfate adenylyltransferase (*cysDN*) (Fig. 1).

### Phylogeny, taxonomic resolution, and average nucleotide identity among chemoautotrophic functional groups

The phylogenetic reconstruction of estuary and intertidal MAGs containing a sufficient number of ribosomal protein encoding genes included 2706 MAGs from the Genomes from Earth’s Microbiomes (GEM) database and 96 recovered in this study. Overall, the tree contained 54 different phyla from 2688 Bacterial and 114 Archaeal genomes. The most species rich phyla included Proteobacteria, Bacteroidota, and Actinobacteriota, which contained 1099, 590, and 274 MAGs respectively (Fig. 3). The MAGs recovered from N-enriched samples were distributed across six of the 54 phyla and included Bacteroidota, Campylobacterota, Deslufuromonadota, Nanoarchaeota, Patescibacteria, and Proteobacteria (Table S3), with 71of the FTR MAGs assigned to Proteobacteria. The unenriched MAGs were distributed across the same phyla, except for Desulfuromonadota (Table S3).

**Figure 3.**
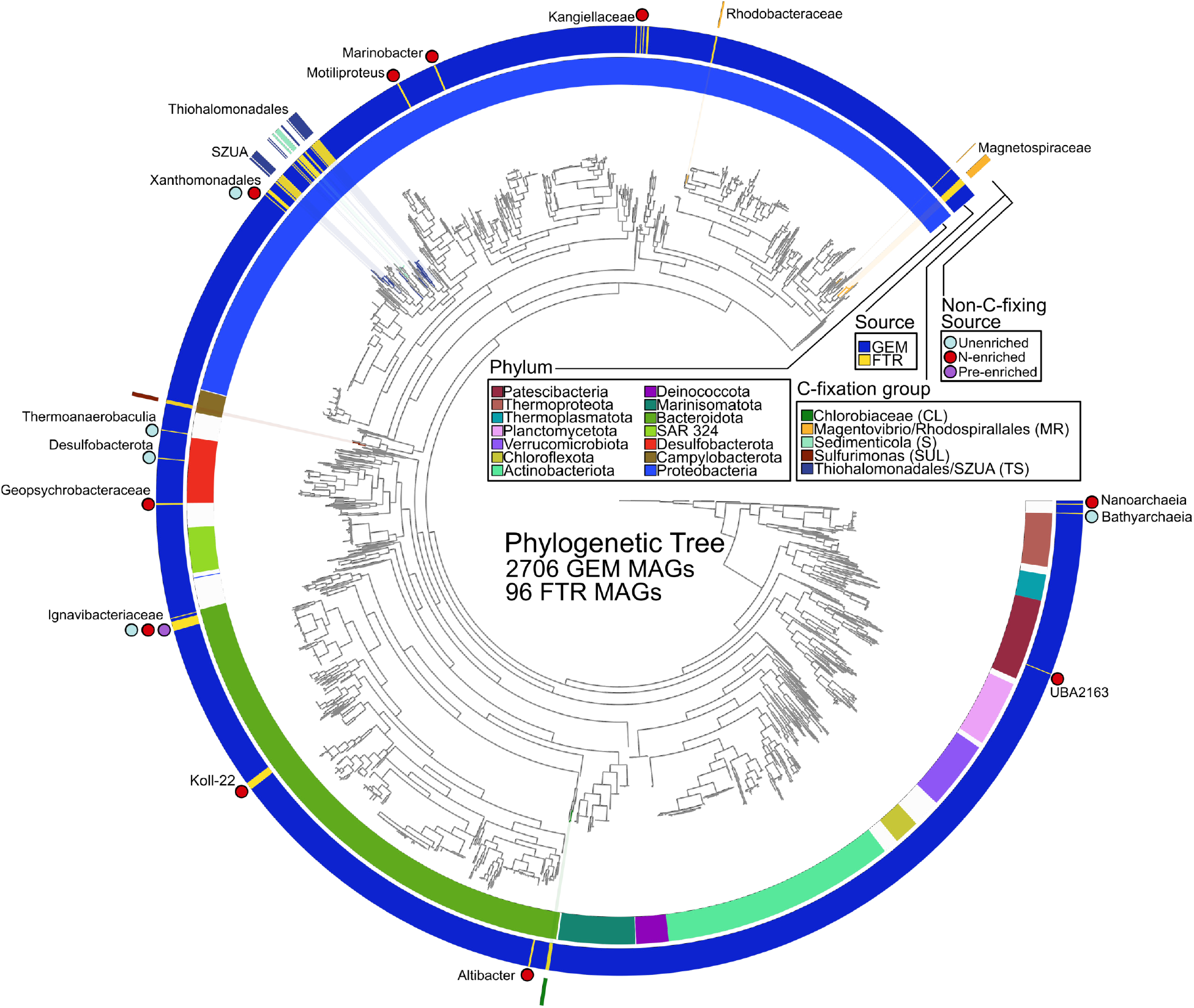
Phylogenetic tree of Genomes from Earth’s Microbiomes (GEM) and the flowthrough reactor (FTR) experiment MAGs. The tree at the center of the display was reconstructed using a minimum of 10 genes in a set of 26 ribosomal protein encoding genes. The first ring around the phylgenetic tree shows the phylum level assignment of each MAG according to the GTDB-Tk. The second ring indicates the source of the MAG (either FTR sediment from this experiment or from marine intertidal or estuary MAGs contained in the GEM database). The third ring shows the FTR C-fixation group assignment of the MAG as bars and the taxonomy of non-C-fixing (non chemoautotrophic) MAGs displayed as text. The circles displayed on the third ring represents the source of the FTR sample.

The carbon fixation groups we identified were distributed across three phyla (Fig. 3). The four non-redundant *Chlorobium* MAGs from the unenriched samples were all located on adjacent leaves within the Bacteroidota and represented the only genomes from this genus in the collection of marine intertidal and estuary samples in our analysis (Fig. 3). They were most closely related to *Chlorobium* recovered from a chemocline in the Black Sea (see pangenomic results). Four of the *Sulfurimonas MAGs* with the potential to fix carbon using the rTCA cycle were found on a single branch among 22 Campylobacterota along with a reference *Sulfurimonas* MAG recovered from the metagenome of inlet seawater (JGI project ID: 3300024257). Three of the *Sulfurimonas* MAGs recovered from mid and deep sediments were redundant. The nine non-redundant *Sedimenticola* were in the same region of the tree as the *Thiohalomonadales/SZUA* group and there were other MAGs from marine systems located on adjacent leaves. There was a clear phylogenetic split of the MAGs classified as Thiohalomonadales and SZUA. The SZUA were unique MAGs whose closest relatives were isolated from a study of hydrothermal vent communities of the Guaymas Basin (34). The Magnetospiraceae and Rhodobacteraceae that formed a cluster based on functional annotation (Fig. 1) were phylogenetically distinct (Fig. 3). The Rhodobacteraceae recovered from the mid and deep depths of N-enriched sediment from core 1 (N1M-Bin_22, N1D-Bin_9) were collapsed into one MAG based on our ANI thresholds (Table S5, Table S6).

The phylogeny of the MAGs that were not classified as chemoautotrophs included eight different phyla (Fig. 3). The Ignavibacteriaceae were recovered from pre-enriched, unenriched, and N-enriched conditions; the Thermoanaerobaculia, Desulfobacterota, and Bathyarchaeia were recovered from unenriched reactors; and several others including Nanoarchaeia, koll-22, and Xanthomonadales were recovered exclusively from N-enriched reactors. The eight MAGs classified as koll-22 were all found to represent a single redundant collection of MAGs and were classified as BRH-c54, a genus level classification assigned to a MAG recovered from the porewater of opalinus clay rock (35). The collection of 10 Ignavibacteriaceae MAGs formed a tight phylogenetic cluster that included reference MAGs from marine wetlands. All Ignavibacteriaceae MAGs were found to be unique non-redundant populations and two of the MAGs (S2M-Bin_6, S2M-Bin_2) were isolated from the same unenriched sample (unenriched core 2). Seven of the Ignavibacteriaceae MAGs were classified as BMS3ABIN03, a genus isolated from sub-seafloor massive sulfide deposits (36). Two MAGs were classified as Archaeae, and one as a member of the candidate phyla Patescibacteria (Fig. 3).

We recovered 97 unique MAGs from the original collection of 113 MAGs according to the threshold of 95% identity over 90% of the length of the MAG (Table S3, Table S5, Table S6).

### MAG relative abundance among N-enriched reactors

Most of the 97 dereplicated MAGs were detected in multiple depths within independent cores, but we observed trends in the abundance that were apparently dependent on depth and N enrichment (Fig. 4). For example, N2S-Bin_4 was detected at each depth in all N-enriched cores, but was not detected in the pre-enriched, unenriched cores, or in-situ samples. This MAG also contained seven other MAGs in the dereplicated cluster (Table S3).

**Figure 4.**
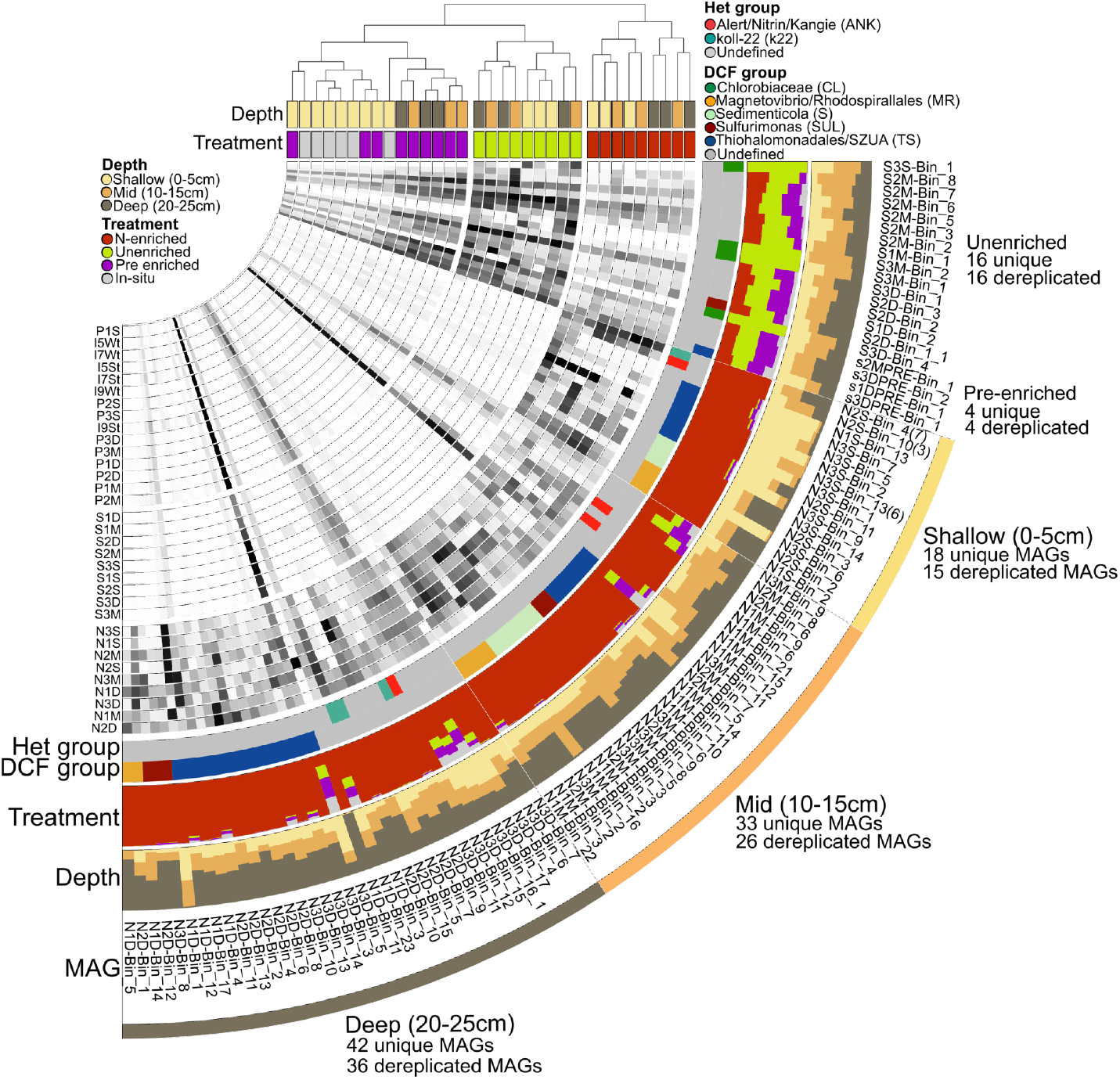
Relative abundance patterns of the 97 non-redundant set of MAGs in each sample. The relative abundance of each MAG is shown as a grey scale heatmap at the root of the display. Values represent log transformed average fold coverage that are also corrected for the number of reads in each sample to allow for comparison among MAGs and across samples. The heterotrophic and dark carbon fixation grouping of each MAG are annotated in the rings layers after the heat map, followed by the proportion of detection in each treatment, proportion of detection at each depth, MAG name, and finally layers indicated the depth where the MAG was recovered. We did not recover MAGs from the in-situ samples. Three of the MAGs show a number in parentheses following the name, this indicates the number of replicate genomes in the cluster. The full taxonomy for each MAG can be found in Table S3. The samples in the heat map are clustered based on Bray-Curtis distances of the corrected relative abundance. Metadata for each sample are provided as color bars beneath the hierarchical tree.

The MAGs recovered from the N-enriched samples were rarely detected in the pre-enriched or unenriched reactors and in-situ samples (Fig. 4). Exceptions to this observation included N3M-Bin_11 and N3D-Bin_5, which are both members of the (TS) group. We also observed that the reads recruited by the N-enriched MAGs were depth dependent. For example, most of the MAGs recovered from the shallow sediments were primarily detected in the shallow samples of our collection, rather than mid or deep samples (Fig. 4). However, MAG N1D-Bin_1 was recovered from a deep fraction but recruited more reads from almost all the other N-enriched samples (Fig. 4). This is likely due to the failure to dereplicate this MAG from N3S-Bin_13. Although N1D-Bin_1 and N3S-Bin_13 shared 99% ANI, the length of the genome alignments was 72% and 49% for each of the pairwise alignments (Table S5, Table S6). The functional similarity (Fig. 1) and taxonomic classification (Table S3) serve as additional lines of evidence that these two MAGs should be considered the same organism, which was recovered from the same core at shallow, mid, and deep fractions. Several additional N-enriched MAGs could also be detected at multiple depth fractions of the same core. For example, N1M-Bin_3 could be detected in three samples (N1S, N1M, and N1D) (Fig. 4).

Apart from the CL group, which were only detected in the unenriched reactor samples, MAGs within the chemoautotrophic functional groups were detected at each depth fraction in all three cores but recovered primarily from the mid and deep depths, where 33 and 42 unique MAGs were recovered respectively (Fig. 4). Members of the chemoautotrophic functional groups were also among the most abundant MAGs in the N-enriched samples (Fig. 4). MAGs within the S functional group were not recovered from the deep sediments and no SUL members were recovered from the shallow sediments (Fig. 4). However, we did detect MAGs representing these groups through read mapping within each of the depths, as shown in Fig. 4. This trend of co-occurrence among MAGs within each functional group was observed within each of the N-enriched samples. The heterotrophic groups were also detected in each of the depths of N-enriched samples. Although the k22 group appears to be absent, members of the dereplicated cluster represented by N2S-Bin_4 were recovered from mid depths (e.g. N3M-Bin_4) (Fig. 1).

### Pangenomics of Chloribium and Sedimenticola

The *Chlorobium* pangenomic analysis included three FTR MAGs and 20 reference genomes classified as *C. phaeobacteroides*, *C. chlorochromatii*, *C. phaeovibriodes*, *C. limicola*, and *C. ferrooxidans*. We identified a total of 5803 gene clusters and 239 core single copy genes. The three FTR *Chlorobium* MAGs formed a cohesive cluster based on phylogenomic and gene cluster content (Fig. S2) and shared the most gene clusters with marine and hot spring derived strains. A *Chlorobium phaeobacteroides* reference genome recovered from a chemocline located 100 m below the surface of the Black sea (GCA_000020545) was most closely related to the FTR *Chlorobium* MAGs according to gene cluster content and phylogeny determined from single copy core genes (Fig S2). The ANI among the FTR *Chlorobium* MAGs and the Black sea reference was above 80%. We identified 443 gene clusters that were unique to the FTR *Chlorobium* MAGs and *C. phaeobacteroides*. Genes within this group encoded proteins related to cofactor and vitamin metabolism, lipid metabolism, ATP synthesis, central carbohydrate metabolism, and a photosystem P840 reaction center cytochrome c551. We also identified a collection of 104 gene clusters that were unique to only the FTR *Chlorobium* MAGs (Fig S2, Table S7). Functions encoded by genes within this cluster included multiple proteins for F-type ATP synthesis and heme biosynthesis.

We chose *Sedimenticola* as the representative genus for MAGs that contain the genetic capacity for the rTCA cycle. The pangenomic analysis of *Sedimenticola* included four FTR MAGs and 14 reference genomes including *S. selenatireducens*, *S. thiotaurini*, and *S. endophacoides*. Pangenomic analysis identified a total of 6811 gene clusters. We identified 262 single copy gene clusters that we used to determine phylogeneomic relationships among genomes. We found that N1M-Bin_16 was most closely related to two *S. selenatireducens* reference genomes (GCA_002868805 and GCA_000428045) isolated from marine sediment (Fig S2). The ANI between N1M-Bin_16 and these genomes was above 88%. The proportion of gene clusters shared with N1M-Bin_16 and GCA_002868805 was highest, but *S. thiotaurini* and other *Sedimenticola* were also all above 80% (Fig. 5). N1M_Bin-2 and N2S_Bin-3 formed a cluster with *S. thiotaurini* (GCA_007713595) and ANI between this reference genome and the FTR MAGs was above 95% in both cases. We identified 220 accessory gene clusters that were found only in FTR MAGs, *S. selenatireducens*, *S. thiotaurini*, and unclassified *Sedimenticola* derived from marine environments. Genes within this cluster encode a wide variety of functions including those related to glyoxolate bypass, glycolysis and rTCA (Table S7). These clusters also contained genes important for cofactor and vitamin metabolism.

**Figure 5.**
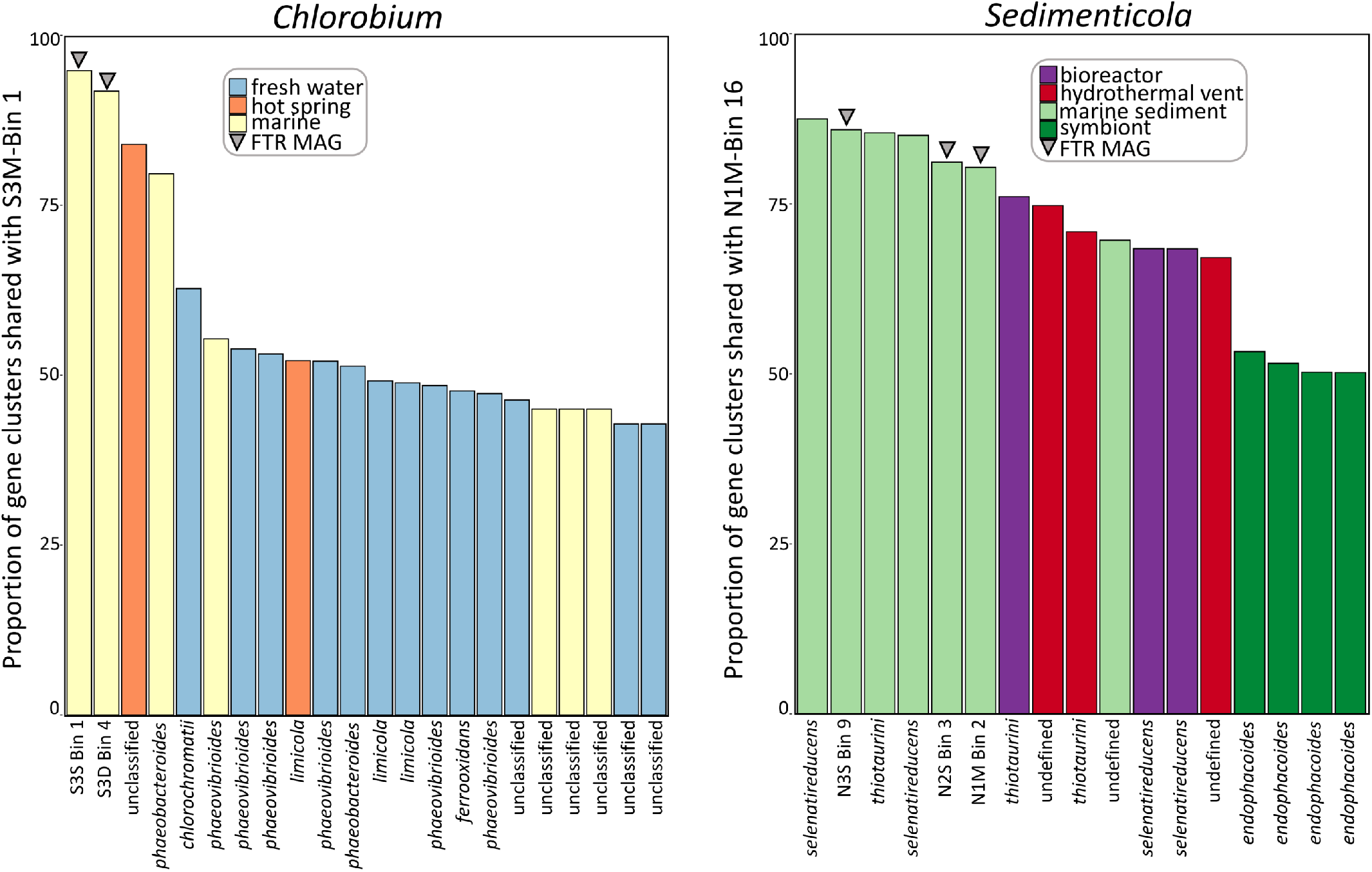
The proportion of gene clusters shared between one of the MAGs recovered in this study with reference genomes and other MAGs recovered within the same genus. The similarity in gene cluster number is based on the assignment of genes into gene clusters according to the *Sedimenticola* and *Chlorobium* pangenomic analysis. The color of the bar indicates the enviroment that the genome was recovered from and the taxonomy of the genome is indicated on the x-axis. An inverted triangle above the bar indicates that the representaitive genome was recovered from metagenomic data in this study.

## Discussion

While the importance of chemoautotrophy in coastal sediments has been recognized for decades (11), few studies have employed genome resolved metagenomic approaches to study them, leaving gaps in our ability to understand biogeochemical processes of the sediment. Our experiment, with high-nitrate, dark, and anoxic conditions applied to sediments for 92 days with no organic matter input, created a niche for chemoautotrophy in each of the three depths examined here. Our observations of chemoautotrophs were stable across three independently collected cores from sediment inhabited by the tall ecotype of *Spartina alternifora*. The reconstruction of 113 MAGs from these sediments enabled us to address three outstanding unknowns about microbial chemoautotrophs in salt marshes, 1) their metabolic flexibility and pathways for DCF, 2) the potential niche space within the top 25 cm of the sediment, and 3) the functional and phylogenetic relationship to microbes inhabiting oligotrophic environments. This study provides much needed genomic insight to improve our understanding of the relevance of DCF to carbon cycling in marine coastal systems and the evolution of DCF within oligotrophic environments more broadly.

We identified three distinct functional strategies for DCF within the FTR cores including CBB driven by denitrifying sulfur oxidation, rTCA driven by denitrifying sulfur oxidation, and rTCA fueled by photoautotrophy despite extremely low light levels (Fig. 1). The use of multiple electron acceptors by the collection of chemoautotrophic MAGs in each of the functional groups could provide an important adaptation to inhabiting the top 25 cm of marsh sediment. Salt marsh sediments contain steep gradients in oxygen (37,38) and nitrate (19,39), and a sulfate to sulfide transition zone (40), all of which are influenced by diurnal tidal inundation that provides a re-supply of fresh organic matter and nutrients (41). The biogeochemistry of the sediment is also influenced by the release of oxygen and carbon from plant roots (42,43) and animal burrows, providing additional structure and permeability to sediment that enhances chemoautotrophic growth (44). These conditions also favor diverse heterotrophic bacteria that primarily rely on sulfate reduction due to the low availability of oxygen and nitrate (45,46). However, the quality of carbon decreases with depth following the mineralization of simple carbon by heterotrophs, reducing the availability of favorable carbon sources used by the heterotrophic community. These conditions present a potential niche space for chemoautotrophic bacteria.

The chemoautotrophic MAGs recovered from the N-enriched reactors contained genes for complete dentification and cytochromes that require oxygen as the terminal electron acceptor among functionally diverse groups of bacteria (Fig. 1, Fig. S3). The ability to use both nitrate and oxygen as terminal electron acceptors, while oxidizing multiple reduced forms of sulfur to produce their own carbon, provides potentially important adaptations to an environment with rapidly changing electron acceptor concentrations and low availability of simple forms of carbon (47). The presence of the different functional groups using the same core metabolism of denitrifying sulfur oxidation suggests that there may be a significant physical niche space allowing for the accumulation of diverse microbes contributing to DCF (48,49).

The presence of these chemoautotrophic groups within the sediment inhabited by the tall ecotype of *S. alterniflora* has important consequences for the ecology of the vegetation and surrounding waters. Many of the chemoautotrophic MAGs are complete denitrifiers (Fig.1, Table S4) and most contain the gene for nitrous oxide reductase (nosZ), which provides the potential to release inert N_2_ from the sediment as opposed to the potent greenhouse gas, N_2_O. The capacity to oxidize reduced sulfur from the sediment is important to the health of *S. alterniflora* due to the toxicity of high sulfide concentrations (50). The carbon storage potential of the sediment could also be affected if appreciable amounts of carbon are being fixed by the chemoautotrophs. Within anoxic sediments below 10 cm, fresh inputs of carbon are likely limited, so the chemoautotrophic necromass may be an important carbon source for the heterotrophic community (51). Cycling of carbon between chemoautotrophic and heterotrophic groups could provide a mechanism to preserve and cycle carbon stored in the system.

The canonical denitrifying MAGs, including members of the *Colwellia*, *Motiliproteus*, and novel genera within the *Kangiellaceae* family, recovered from the N-enriched sediment were largely complete denitrifiers (Fig. 1). However, we observed the carbon sources potentially used by most heterotrophic denitrifiers were the same as the chemoautotrophic denitrifying sulfur oxidizers, including chitin and polyphenolics. Three *Colwellia* redundant MAGs recovered from three different depths and two separate cores contained evidence for more extensive use of plant derived carbon, primarily from the decomposition of hemicellulose (Fig. S3). The lack of a significant carbohydrate usage profile among most of the heterotrophic denitrifiers (Fig. 1, Fig. S3) provides evidence that they may rely on more simple forms of carbon derived from the chemoautotrophic necromass or through syntrophic interactions. Coupling between chemoautotrophic carbon fixation fueling heterotrophic communities can be found around hydrothermal vents (52), and in benthic freshwater ecosystems (53). Recent evidence also suggests that low abundance chemoautotrophic members of the community can support heterotrophic communities within deep sea oligotrophic sediment through syntrophic interactions and metabolic handoffs (54). Quantifying the flow of organic carbon among the heterotrophs and autotrophs in the sediment will provide important insights about the flow and retention of carbon within these blue carbon systems.

While chemoautotrophs may be low abundance members of the unenriched community (Fig. 4), the findings here suggest that there may be conditions in the heterogeneous vegetated salt marsh platform where they are abundant. The relative abundance estimates, and average nucleotide identity (ANI) of MAGs obtained from the three independent depth profiles indicates that some populations of chemoautotrophs spanned more than one depth fraction of the vertical profile and/or multiple cores (Fig.4, Table S3). Support for the presence of the same population represented by an individual MAG includes 1) the observation that redundant genomes were reconstructed within two or more N-enriched sample cores and/or depths and 2) read mapping that indicates detection of the same MAG within multiple samples (Fig. 4). Although we did not have the required coverage for a detailed strain analysis (55), the level of nucleotide similarity (95%) and length (90%) thresholds used to evaluate redundancy among our collection of MAGs is strong support for the occurrence of highly similar genomes. For example, two *Thiohalomonadaceae* MAGs, recovered from shallow and mid-depth sediments contained 99.9% ANI at the nucleotide level over 98.8% of the genome (Table S5). *Sulfurimonas* MAGs were also recovered from multiple cores and contained similar alignment length percentages and ANI. These findings suggest that some populations may be large and capable of inhabiting different niches within the sediment. *Sulfurimonas* is known to contain organisms that are metabolically plastic and ubiquitous across the globe (56). Additional sequencing of the N-enriched reactor samples or additional enrichment experiments would be useful to identifying the strain level variation of these communities.

Gammaproteobacteria are widespread and active members of the microbial communities inhabiting marine sediments (10) and the MAGs recovered from the N-enriched reactors expands our understanding of the functional metabolism that may be important to their ubiquitous nature. The experimental nitrate amendment allowed us to recover chemoautotrophic members of this phylum that were undetectable in the unenriched FTR or in-situ samples but represented an average estimate of 13% of the total community within nitrate enriched sediment. Our evaluation of the phylogenetic breadth of environmental genomes recovered from intertidal marine sediments identified 54 different phyla and six of these phyla are represented by MAGs in the N-enriched reactors (Fig. 3). This points to a relatively narrow phylogenetic group that are enriched by these conditions and this finding is confirmed by the reduced alpha diversity estimates for these samples observed by Bulseco *et al*. (19). Except for *Chlorobium*, the presence of close phylogenetic relatives within coastal marine systems indicates that the MAGs recovered here are not necessarily limited to salt marsh habitats. The chemoautotrophic MAGs in this study were distributed among 12 different families and many of these families have evolutionary relationships to metabolically diverse microbes inhabiting oligotrophic environments. For example, the families identified as SZUA-229 and SZUA-152 are Gammaproteobacteria originally recovered from oxic subseafloor sediments adjacent to a cold water aquifer (57). The *Sulfurimonas* potentially conduct DCF in salt marsh sediments (58), deep sea hydrothermal vents (59), and pelagic redox clines (60). The *Sedimenticola* are related to inhabitants of other oligotrophic environments including hydrothermal vents (61) and marine sediment (62). Unenriched FTR sediments also contained *Chlorobium*, closely related to another species inhabiting a chemocline in the Black Sea (63,64). While the taxonomic relationship among the chemoautotrophic MAGs in this study is consistent with ubiquitous chemoautotrophic Gammaproteobacteria in marine systems, this study improves our ability to understand their underlying functional strategies.

To identify genomic features that are important for inhabiting such diverse environments, we conducted pangenomic analysis of the *Chlorobium* and *Sedimenticola* MAGs. Gene clusters shared among the *Chlorobium* MAGs recovered here with the *C. phaeobacteroides* strain recovered from the Black Sea, included a P840 reaction center and an F-type ATPase, which harvest light and provide proton regulation for the cell, respectively. We also identified unique gene clusters orthologous to vitamin and cofactor metabolism, glycan and lipid metabolism, and energy production (Table S7). These functional categories can be used to define differences between oligotrophic and eutrophic environments (65) and are important to determining the flow of carbon between soil organic matter and the microbial community (66). Oligotrophic conditions and a strong redox cline are two defining characteristics of the Black Sea chemocline as well as the sediment in our FTR experiment (19,67), suggesting that these are two important characteristics of the *Chlorobium* niche space.

Pangenomics of *Sedimenticola* genomes revealed a cluster of genes shared among all *Sedimenticola* recovered from marine environments. The cluster contains genes related to carbohydrate and amino acid metabolism, metabolism of cofactors and vitamins, and aromatics degradation (Table S7). Strains of *Sedimenticola* recovered from salt marshes show improved growth in the presence of amino acids (62) while the originally described strain, *Sedimenticola selenatireduces*, couples the oxidation of aromatic compounds to selenate respiration (68). Interestingly, the *Sedimenticola* recovered from the FTR experiment shared the most gene clusters in the pangenome with *S. selenatireducens*. These findings highlight the complexity of microbial carbon cycling that can be recovered using genome resolved metagenomics and presents many questions about the nature of the microbial taxa recovered from N-enriched systems.

The MAGs recovered from these nitrate enriched FTRs improve our understanding of DCF within vegetated coastal marine systems and other oligotrophic environments. Our research expands the known metabolic flexibility among Gammaproteobacteria inhabiting marine and oligotrophic sediments, extends our understanding of the niche space and population sizes of chemoautotrophs, and provides the phylogenetic context of members of the microbial community enriched by nitrate. Few chemoautotrophic genomes have been recovered from metagenomic datasets and identification of a significant niche space within salt marsh sediments will renew interest in understanding their contribution to nitrogen, sulfur, and carbon cycling. The potential for using oxygen and nitrate as a terminal electron acceptor and the evidence for chitin mineralization requires further investigation, but the observation of this metabolic potential across multiple chemoautotrophic functional groups suggests that chemoautotrophy and/or mixotrophy may be an important feature of microbial life in nitrate enriched sediments. In-situ biogeochemical measurements of chemoautotrophy paired with targeted cultivation and enrichment represent two complimentary methodologies that should be used to assess DCF relevance in these critically important coastal carbon storage systems.

## Acknowledgements

We acknowledge the Bay Paul Center at the Marine Biological Laboratory for sequencing support. Funding or this project was provided by an NSF CAREER Grant to JLB (DEB1350491) and an NSF DDIG to JLB and ANB (Award # 1701748). The original FTR experiments were supported by a Woods Hole Sea Grant award (Project # NA140AR4170074 Project R/M-65s to Anne Giblin) and a Ford Foundation pre-doctoral fellowship to ANB. Samples were collected from the Plum Island Ecosystems LTER, which is supported by NSF (OCE 0423565, 1058747, and 1637630) and from the NSF funded TIDE project (DEB 1902712). Samples from this project were processed with equipment purchased under an NSF FMSL grant to Northeastern University (DBI 1722553). This publication is number TBD from the Marine Science Center.

